# Prenatal cannabis smoke exposure alters placental development in a murine model of pregnancy

**DOI:** 10.1101/2025.06.28.662155

**Authors:** Tina Podinic, Maria Sunil, Andie MacAndrew, Cristina Monaco, Grace Lee, Cielle Lockington, Jim Petrik, Amica-Maria Lucas, Thane Tomy, Gregg Tomy, Joanna Kasinska, Laiba Jamshed, Alison C. Holloway, Elyanne M. Ratcliffe, Sandeep Raha

## Abstract

Cannabis use during pregnancy continues to increase with smoking remaining the most common mode of consumption. While clinical studies highlight an association between prenatal cannabis use and adverse pregnancy outcomes, less is known about placental outcomes, even though many of the reported pregnancy outcomes are thought to be mediated via placental dysfunction. Here, we established a mouse model of gestational cannabis smoke exposure to investigate the impacts on fetal outcomes and placental structure and function. Pregnant CD1 mice were exposed daily to Δ9-tetrahydrocannabinol (THC)-dominant cannabis smoke (12-14% THC, 0-2% CBD) or filtered air from embryonic day (E)6.5 to E18.5 or parturition. Cannabinoid analyses in cannabis smoke-exposed, paired maternal and fetal livers revealed total THC and 11-Nor-9-carboxy-THC (THCA) concentrations of 135.95 ± 13.60 ng/g and 30.84 ± 4.68 ng/g, respectively. Moreover, *Cyp1a1*, a smoke-inducible enzyme, was induced by 4-fold in cannabis smoke-exposed placentae. No changes in offspring body weights were observed; however, there was a marked decrease in the brain-to-body weight ratio of exposed postnatal day 1 (PND1) offspring. Placentae from exposed dams were significantly reduced in size, with altered zonation marked by a significantly decreased junctional zone and increased labyrinth zone. Key trophoblast differentiation markers (*Tfap2c*, *Tpbpa*, *Pcdh12*) and placental endocrine regulators (*Pl2*, *Igf1r*) were significantly downregulated following cannabis smoke exposure in placentas. Furthermore, transcript levels of placental nutrient and vascularization markers, *Glut1*, *Vegfa* and *Pparg* were significantly decreased in cannabis smoke-exposed placentas. By employing a physiologically relevant platform of prenatal cannabis exposure in vivo we demonstrate the adverse effects of prenatal cannabis smoke exposure on placental structure and function as well as on fetal brain growth.

## Introduction

Cannabis use during pregnancy is becoming increasingly prevalent, with 2-20% of pregnant women reporting prenatal cannabis use for relief from common pregnancy symptoms, such as anxiety, pain and nausea, as well as for recreational purposes (1–8). In recent years, cannabis products have evolved to include formulations like vapes, edibles and oils; however, most pregnant women still report smoking cannabis (9, 10). Although human studies have shown that in utero cannabis exposure is associated with low birthweight (11, 12), elevated risk of fetal death (11), small-for-gestational age (SGA), preterm birth (12) and adverse neurodevelopmental outcomes in offspring (13), there is still conflicting evidence due to confounding variables, such as the use of multiple substances, variations in cannabis potency and frequency of use.

*Cannabis* (marijuana) is composed of over 550 compounds. The phytocannabinoids, including Δ9-tetrahydrocannabinol (THC) and cannabidiol (CBD), comprise approximately one-fifth of these chemical constituents (14). In addition to phytocannabinoids, marijuana contains several other bioactive components, including terpenes, flavonoids and phenolic compounds. Despite this chemical complexity, existing preclinical studies of cannabis exposure in pregnancy have primarily employed single-component analyses to investigate the biological effects of THC (15–17) or CBD (18, 19), as well as their closely related metabolites. There are few studies that interrogate the contribution of the remaining components and combustion byproducts associated with smoking cannabis. Like tobacco, cannabis smoke contains polycyclic aromatic hydrocarbons (PAHs), whose contributions may be overlooked in studies of their bioactive components, THC and CBD (20, 21). Furthermore, the mode of administration of cannabis, whether through inhalation (smoking, vaping) or ingestion (edibles, oils), yield distinct pharmacokinetic and pharmacodynamic profiles, ultimately altering the overall concentrations, tissue distribution and duration of actions of individual cannabinoids or their associated metabolites (22–24). To add another layer of complexity, phytocannabinoids and terpenes possess synergistic and/or antagonistic effects through their overlapping interactions with receptors belonging to the endocannabinoid system (ECS), also referred to as the “entourage effect” (25).

Increasingly, studies are interrogating the complex effects of cannabis on aspects of female reproduction, including fertilization, implantation and placentation (26–28). Being the primary maternal-fetal interface, the placenta is responsible for supplying the developing fetus with nutrients and growth hormones, facilitating gas and waste exchange, and providing structural protection by limiting the fetus’ exposure to toxicants (29). As such, the placenta plays a significant role in fetal development and health (30, 31), emphasizing the importance in understanding the impact of the various components of cannabis on this organ. Prenatal THC exposure in preclinical models has been associated with placental insufficiency (17), a condition characterized by defects in placental vascular remodeling leading to fetal hypoxemia (17). More specifically, placental insufficiency commonly resulting from abnormal placental vascularization, refers to the inability of the placenta to deliver nutrients and oxygen to the fetus. This effect of THC to impair placental function is consistent with reports of fetal growth restriction (FGR) following THC exposure (16).

While studies have reported placenta-specific effects of THC exposure, to date, few studies have investigated the impacts of gestational cannabis smoke exposure in vivo (23, 32). In a 2017 study, Benevenuto and colleagues employed a cannabis smoke exposure model designed to mimic recreational use during pregnancy. These authors observed enlarged placental size, suggesting that inhaled cannabis may interfere with placentation (33); however, the nature of the changes in the placenta were not reported. Therefore, the impact of prenatal cannabis smoke exposure on placental function and development remains a gap in the current literature. Here, we report that THC-dominant cannabis smoke exposure during pregnancy impacts the structure of placentae and results in reduced brain weight at birth. Furthermore, we confirmed the transmission of THC to fetal tissues following prenatal cannabis smoke exposure and observed altered markers of placental vascularization and altered expression of critical placental hormones.

## Materials and Methods

### Animal model

All animal work was approved by the McMaster University Animal Research Ethic Board under Animal Utilization Protocol number 21-06-16. Six week-old CD1 virgin female mice were purchased from Charles River (Quebec, Canada) and allowed to acclimatize for 2 weeks under standard conditions: 12-h light and dark cycle with a high-fat diet (22% kcal fat, 55% kcal carbohydrates, 23% kcal protein, 3.3 kcal/g; Teklad Global 19% Protein Extruded Rodent Diet, 2919) and water ad libitum at the McMaster University Central Animal Facility. All procedures were approved by the Animal Ethics Research Board at McMaster University (Animal Utilization Protocol 21-06-16). Female mice were harem bred overnight and the presence of a vaginal plug the following day marked embryonic day 0.5 (E0.5). Experimental groups of pregnant mice were randomized into two experimental groups: (1) sham-exposed (CON; filtered air) or (2) cannabis-exposed (EXP; whole cannabis smoke). Pregnant dams were individually housed and subjected to daily exposures to filtered air or cannabis smoke. Dam food consumption and body weight was measured every four days until E18.5 or parturition. A subset of pregnant dams were sacrificed at E18.5 for fetal and placental collections, and the remainder of the dams were allowed to deliver normally. Following parturition (PND1), pups were counted, weighed and the brain weight was determined.

### Cannabis cigarette preparation and smoke exposure protocol

Cannabis cigarettes were prepared as described previously (34) with the Indica-dominant hybrid cannabis strain, Pineapple Kush (12-14% THC: 0-2% CBD) purchased from Cannalogue (Toronto, Canada). 0.85g dry cannabis flower was weighed and packed into empty king-size cigarette tubes. Filters were removed before cannabis cigarettes were loaded into the Smoke Inhalation Unit 24 (SIU 24) system. Pregnant mice were placed into a designated cage within the chamber for the duration of exposure. Six cannabis cigarettes were used in each exposure for an approximate exposure duration of 30 minutes. The sham cohort was placed in an identical and separate exposure cage and exposed to filtered air for the same duration to minimize the variables of handling and confinement stress.

### Analysis of outcomes in E18.5 offspring

Pregnant mice were exposed to filtered air (control: N=9 dams) or first-hand cannabis cigarette smoke (cannabis-exposed: N=10 dams) for a single 30-minute period daily from E6.5 up to E18.5, at which point they were euthanized for fetal tissue collection(s). At euthanasia we collected maternal and fetal blood, maternal and fetal livers, and placentae. Placentae were either snap-frozen in liquid N_2_ or preserved in 4% PFA (Millipore Sigma, 100496) for 48 hours and subsequently embedded in paraffin, for biochemical or immunohistochemical analyses, respectively. To confirm fetal exposure to the major constituents of cannabis, paired maternal and fetal liver samples were collected from cannabis-exposed animals and targeted analyses for THC, CBD and THCA were performed by high pressure liquid chromatography tandem mass spectrometry (HPLC-MS/MS).

### Immunohistochemistry

Paraformaldehyde (PFA)-fixed and paraffin-embedded E18.5 placentae were cut into 5µm sections. Tissues were deparaffinized in xylene and incubated in graded ethanol to rehydrate tissues. To determine the area of the junctional zone of the placentae, Periodic acid-Schiff (PAS) staining was performed by the Histology Core at the McMaster Immunology Research Center (MIRC) using a standard PAS special stain, and imaged on brightfield using a Nikon Ti2 Eclipse microscope. Areas of junctional and labyrinth zones were quantified in CON (n = 3) and EXP (n = 6) placental sections (1 section/fetus/dam) using the Nikon Ti2 Analysis Software and expressed as a ratio over total placental area x 100 (%). Significance was determined via a Welch’s t-test.

To assess if cannabis smoke exposure resulted in placental hypoxia, protein expression of carbonic anhydrase (CAIX) was visualized by immunostaining. Endogenous peroxidase activity was quenched with 3% hydrogen peroxide for 10 minutes, followed by antigen retrieval using citrate buffer with Tween20 (Millipore Sigma, P1379). To reduce non-specific binding of antibodies, samples were blocked in 5% bovine serum albumin (BSA; Millipore Sigma, A8806) with 0.02% sodium azide for 10 minutes at room temperature. Placental sections were incubated in a polyclonal goat anti-carbonic anhydrase antibody (R&DSystems, AF2344, 1:200 dilution) overnight at 4°C. Biotin-conjugated rabbit anti-goat secondary antibody (Abcam, ab6740, 1:100 dilution) was added for 2 hours at room temperature, followed by treatment with ExtrAvidin-Peroxidase (Sigma-Aldrich, E2886, 1:50) for 1 hour. Tissues were exposed to SigmaFast 3,30-diaminobenzidine (DAB; Sigma-Aldrich, D4293) for visualization of staining and counterstained using Carazzi’s hematoxylin. Immunostaining was quantified using morphometry software (Image J, version 1.54p). Total positive CAIX staining in CON (n = 3) or EXP (n = 6) was ratioed with total placental area in each placental section and expressed as a percentage (%) of total placental area (2 sections/placenta/dam). Significance was determined via a Welch’s t-test. All antibodies used in this study are listed in Table S1.

### RNA Extraction and quantitative PCR

50 mg placental tissues were mechanically homogenized in 500 μL ice-cold TRI-zol^TM^ reagent (ThermoFisher, 15596026), followed by a 10-minute centrifugation at 12,000 x g. Following this process, RNA was isolated and filtered using the Direct-zol^TM^ RNA extraction kit (Zymo, R2060) according to the manufacturer’s protocol. RNA concentrations (ng/μl) and purity values were determined using the Nanodrop One^TM^ spectrophotometry instrument (ThermoFisher, ND-ONE-W). 1 μg of RNA was converted to cDNA with the Applied Biosystems^TM^ High-Capacity cDNA Reverse Transcription Kit (Applied Biosystems, 4368813) using the manufacturer’s protocol. RT-qPCR was performed using the Bio-Rad CFX384 Touch^TM^ Real-Time PCR Detection System using GB-Amp™ InFluor^TM^ qPCR Mix (GeneBio, P2092) master mix. Analysis of qPCR results were carried out using the ΔΔCt method (35) and normalized to the geomean of two housekeeping genes (*Actb*, *Rn18s*). All primers used in this study are listed in Table S2.

### Quantification of Cannabinoids in Liver

Unlabeled and mass-labeled analytical standards of exo-cannabinoids, Δ9-THC, CBD and THC-A (>98%) (11-nor-9-carboxy THC), the predominant secondary metabolite of Δ^9^-THC and a proxy for Δ9-THC exposure, were purchased from Toronto Research Chemicals (Ontario, Canada) and utilized as references for targeted analysis of cannabinoids. High performance liquid chromatography tandem mass spectrometry was performed on mouse liver (≥20 mg) and quantified for cannabinoids using an Agilent 1100 with a CTC PAL autosampler with a Sciex 365 triple quadrupole mass spectrometer modified with an HSID Ionics Eclipse Plus orthogonal ionization source. Details of the extraction and multiple reaction monitoring ion transitions for all analytes and mass labelled internal standards have been previously described (36).

### Statistical Analyses

Statistical analyses were performed using GraphPad Prism software V6.0. Normality was assessed for all datasets. Parametric tests were used for normally distributed data and non-parametric Mann-Whitney test were used for non-normal data. A data set with two groups was compared using a two-tailed Student’s t-test, whereas data sets containing two or more groups were compared with a one or two-way analysis of variance (ANOVA) and corrected by a Tukey multiple comparison test. Where there were unequal sample sizes between comparison groups, a Welch’s correction was incorporated. Data are reported as mean ± SEM. Post hoc significance (p < 0.05) will be denoted as bolded numerical p-values.

## Results

### Maternal exposure to cannabis smoke results in transplacental transfer of THC and induction of placental xenobiotic metabolism

Pregnant mice were exposed to cannabis smoke (12-14% THC; 0-2% CBD) daily from embryonic day 6.5 (E6.5) through E18.5 or parturition (**Fig. 1A**). We measured maternal weight gain and food intake, and estimated daily energy intake throughout the exposure window. There were no significant differences between control and cannabis smoke-exposed dams (**Fig. S1**). To estimate trans-placental passage of THC to the fetus, we assessed the total levels of Δ^9^-THC and THCA in paired cannabis-exposed maternal and fetal livers via LC-MS/MS. We determined the cumulative Δ^9^-THC and THC-A levels in maternal and fetal livers to be 135.95 ± 13.60 ng/g and 30.84 ± 4.68 ng/g, respectively (**Fig. 1B**), whereas CBD levels ranged from 4.11-22.68 ± 1.86 ng/g in maternal livers and 3.78-37.13 ± 3.34 ng/g in fetal livers following cannabis smoke exposure. In addition, the expression of *Cyp1a1,* whose gene product is a cytochrome-P450 enzyme potently induced by smoke, was significantly upregulated by 4-fold in cannabis-exposed placentae (**Fig. 1C**).

**Figure 1.**
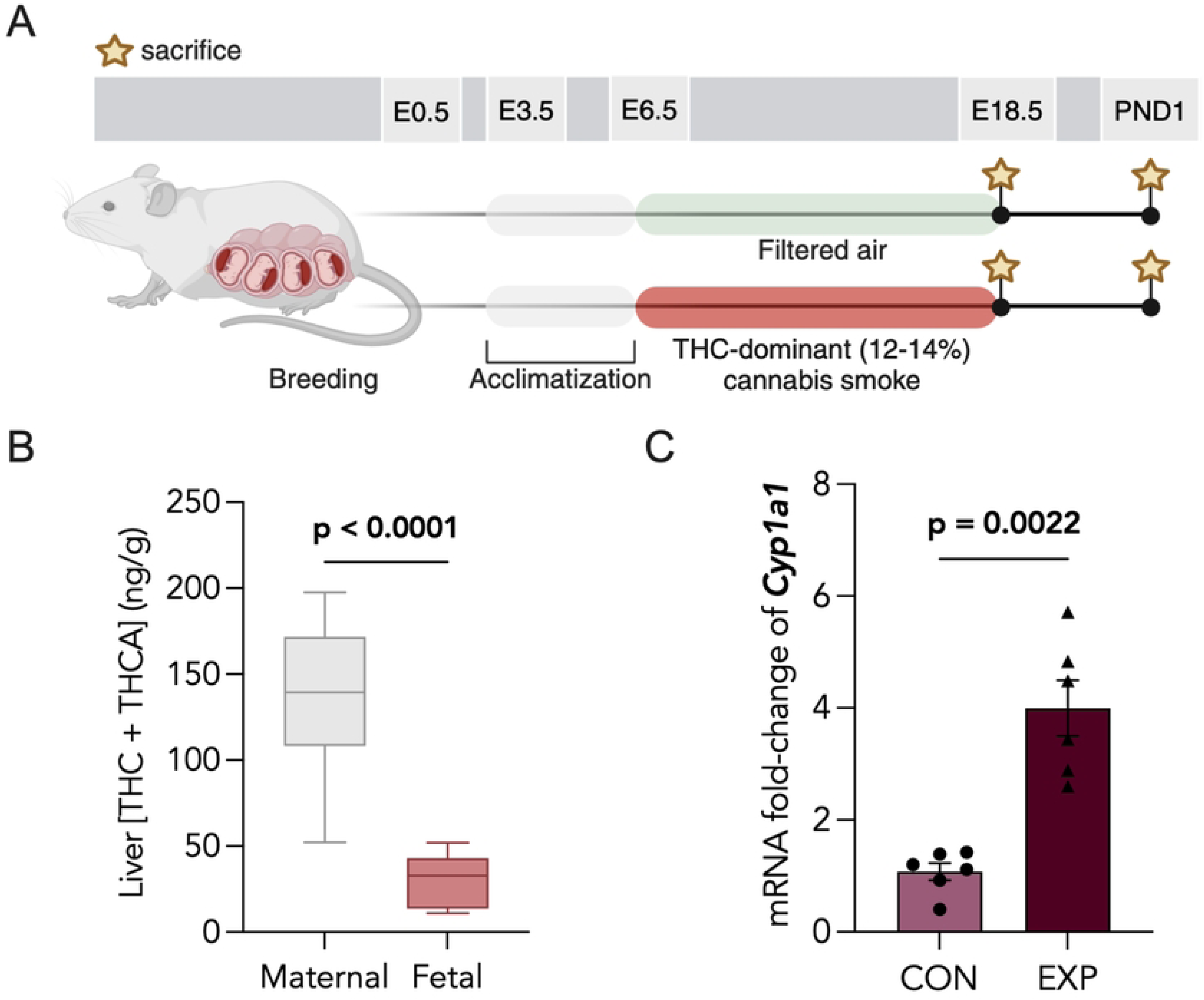
Prenatal cannabis smoke exposure leads to fetal hepatic cannabinoid detection and placental Cyp1a1 induction. (**A**) Schematic of experimental paradigm for prenatal cannabis smoke exposure. Pregnant CD1 mice were exposed to filtered air (CON) or cannabis smoke (EXP; 12-14% THC and 0-2% CBD) (EXP) daily from embryonic day 6.5 (6.5) until E18.5 or parturition. Maternal and offspring tissues were collected at either E18.5 or postnatal day 1 (PND1). (**B**) LC-MS/MS quantification of total THC and THCA in cannabis-exposed livers of dams (n = 10) and corresponding fetal livers (n = 10), in ng/g. (**D**) Gene expression level of *Cyp1a1* was assessed in CON and EXP placentas using RT-qPCR and normalized to the GEOMEAN of two housekeepers, *Actb* and *Rn18s*. Data points represent the mean ± SEM of 6 biological replicates. Statistical significance is denoted by bolded p-values.

### Cannabis smoke exposure leads to increased fetal-to-placental weight ratios and decreased PND1 brain-to-body weight ratios

No significant differences were observed in litter sizes following prenatal cannabis exposure (**Fig. 2A**). Furthermore, no statistically significant differences in bodyweights were observed in E18.5 or PND1 pups (**Fig. 2B, 2C**). Interestingly, cannabis-exposed placentae were significantly smaller at E18.5 (**Fig. 2D**). While offspring body weights were not significantly altered by cannabis exposure, cannabis-exposed PND1 offspring exhibited decreased brain-to-body weight ratios, a difference that was not observed at E18.5 (**Fig. 2E-F**). The fetal-to-placental (F:P) weight ratio was assessed (37). This ratio significantly increased in fetal-placental units exposed to cannabis smoke (**Fig. 2G**).

**Figure 2.**
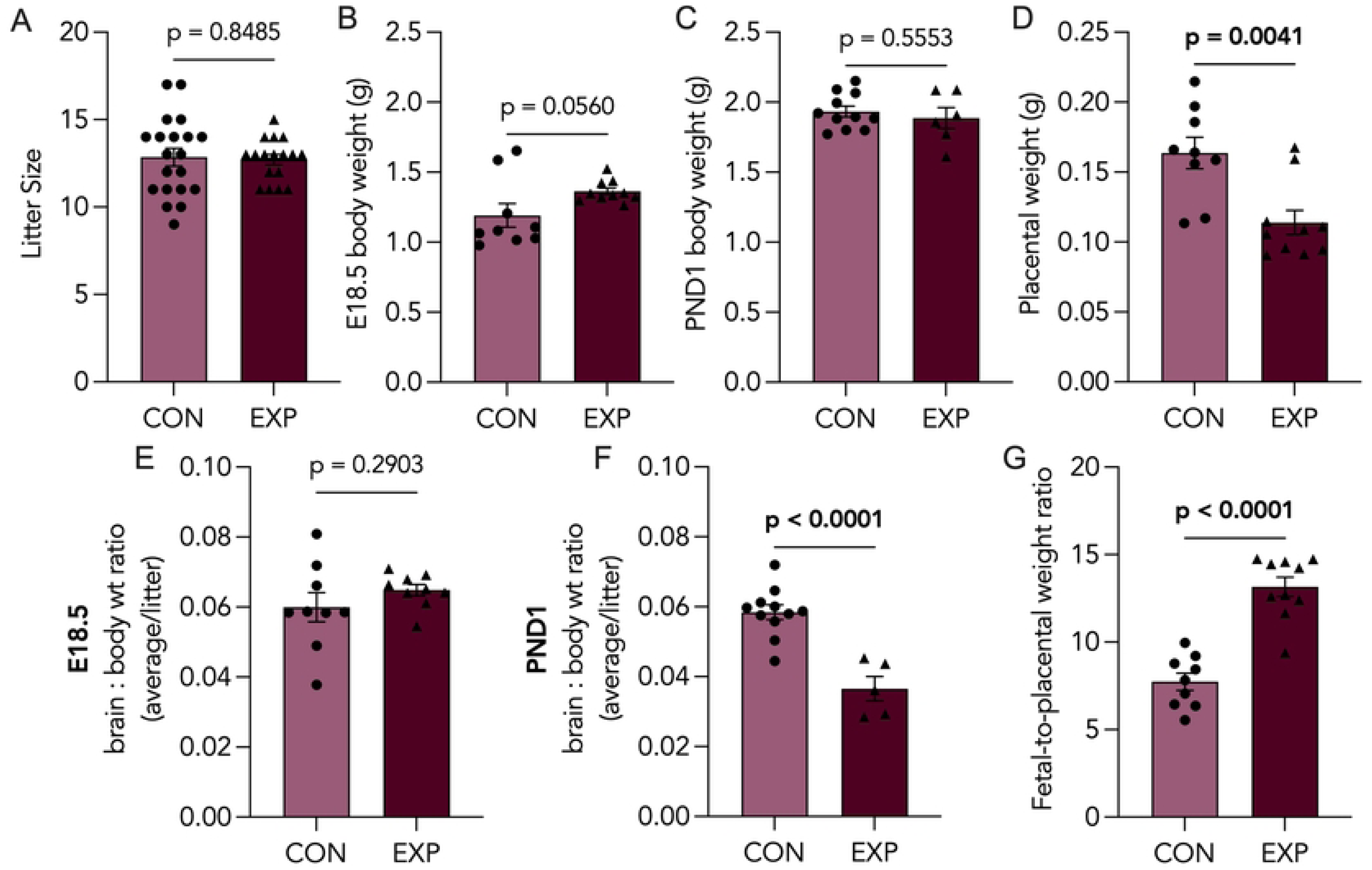
**Biometric outcomes associated with prenatal cannabis smoke exposure**. (**A**) Litter sizes of control (CON; n = 20) and cannabis-exposed (EXP; n = 17) groups from both embryonic day 18.5 (E18.5) and postnatal day 1 (PND1) cohorts. Body weights of (**B**) E18.5 or (**C**) PND1 offspring from CON (n = 9-11) and EXP (n ≥ 6) groups, represented as an average per litter, in grams. (**D**) Placental weights from CON (n = 9) and EXP (n = 10) groups, represented as an average per litter, in grams. Brain-to-body weight ratios of CON (n *≥* 9) and EXP (n *≥* 6) offspring at (**E**) E18.5 and (**F**) PND1, represented as an average per litter. (**G**) fetal-to-placental weight ratio of CON (n = 9) and EXP (n = 10) groups at E18.5, represented as an average per litter. Significant differences are denoted by bolded p-values.

### Placental zonation, developmental and endocrine markers are altered following cannabis smoke exposure

Considering our observation of smaller placental size, we assessed whether there were differences in placental zonation. We quantified the areas of junctional (Jz) and labyrinth (Lz) zones from PAS-stained placental sections (Fig. 3A), a stain commonly used to identify the glycogen-rich Jz (38). We found the percentage of the Jz area over total placental area to be significantly reduced (Fig. 3B), whereas the Lz area was significantly increased (Fig. 3C). Next, we sought to characterize the levels of key placental developmental markers in late pregnancy, transcription factor AP-2 gamma (*Tfap2c*), trophoblast specific protein alpha (*Tpbpa*) and pro-cadherin 12 (*Pcdh12*), which were all downregulated in cannabis-exposed placentae (Fig. 3D**-F**). Due to the decreased area of the junctional zone, which represents the primary endocrine layer, we further assessed the levels of placental lactogen 2 (*Pl2*) and insulin-like growth factor 1 receptor (*Igf1r*), which were significantly downregulated in cannabis-smoke exposed placentae compared to control (Fig. 4A**, 4B**).

**Figure 3.**
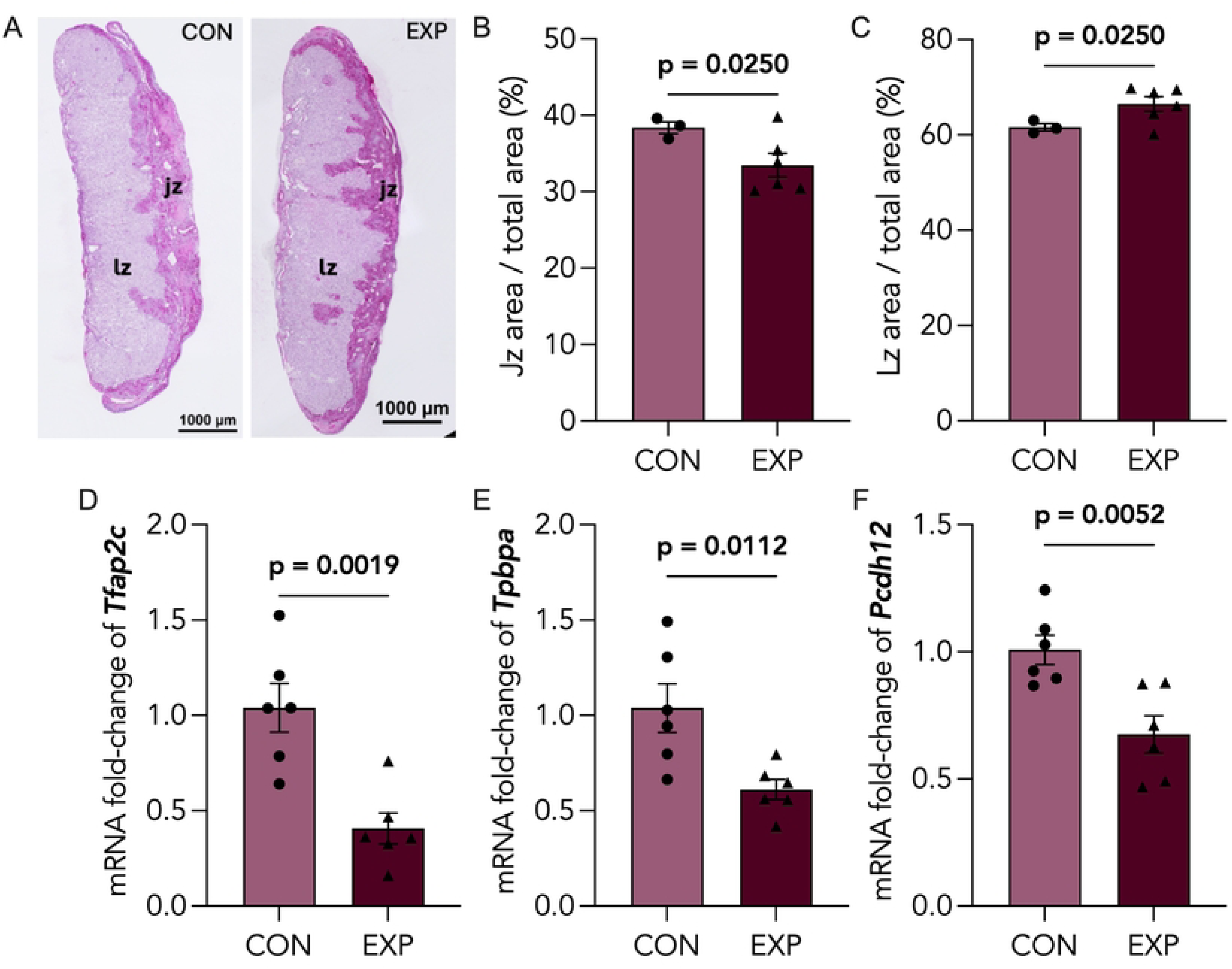
Cannabis-exposed placentas exhibit decreased expression of placental developmental markers and altered zonation. (**A**) Representative image of control (CON; n=3) and cannabis-exposed (EXP; n=6) placentas stained with Periodic acid-Schiff (PAS) stain and with junctional, jz, and labyrinth, lz, zones indicated. Scale bars = 1000µm. Quantification of (**B**) Jz area over total placental area and (**C**) Lz area over total placental area, represented as a percentage. Data points represent the mean ± SEM of 3-6 biological replicates. Significant differences were determined by a Welch’s t-test: *P < 0.05; ****P < 0.0001 relative to control. CON and EXP placentas were assessed for mRNA expression of (**D**) *Tfap2c*, (**E**) *Tpbpa*, and (**F**) *Pcdh12* via RT-qPCR, and normalized to *Actb* and *Rn18s.* Data points represent the mean ± SEM of >5 biological replicates. Significant differences are denoted by bolded p-values.

**Figure 4.**
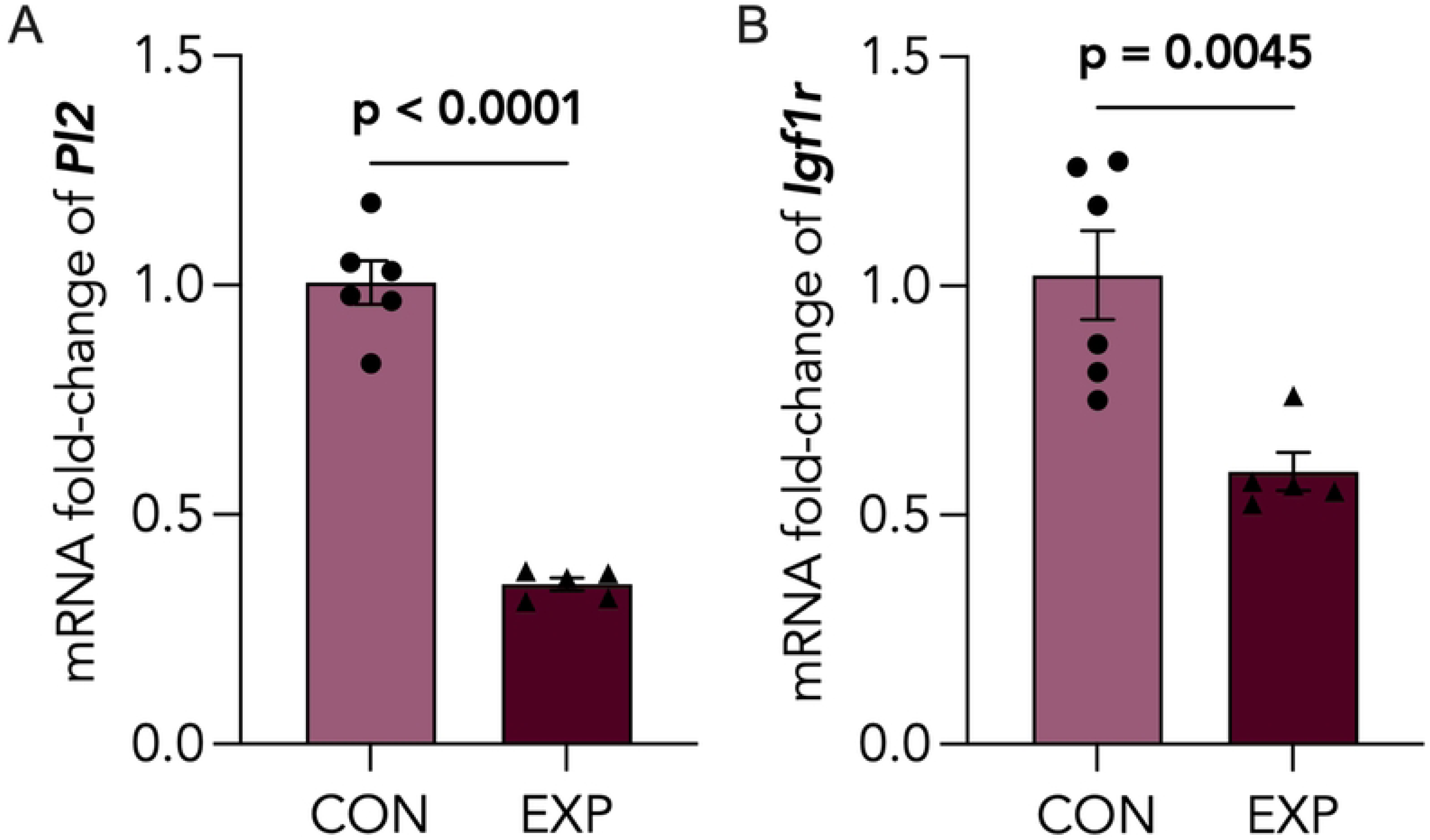
Decreased expression of placental Pl2 and Igf1r following cannabis smoke exposure. Control (CON) and cannabis-exposed (EXP) placentas were assessed for mRNA expression of (**A**) *Pl2* and (**B**) *Igf1r* via RT-qPCR, and normalized to *Actb* and *Rn18s.* Data points represent the mean ± SEM of >5 biological replicates. Significant differences are denoted by bolded p-values.

### Placental nutrient transport and vascularization markers are decreased following prenatal cannabis exposure

To examine the impact of cannabis exposure on placental nutrient transport and vascularization, we assessed mRNA levels of glucose transporter 1 (*Glut1*), vascular endothelial growth factor 1 (*Vegfa*) and peroxisome proliferator-activated receptor γ (*Pparg*). We found that levels of placental *Glut1*, *Vegfa* and *Pparg* are significantly decreased following cannabis exposure (Fig. 5A**-C**). To further assess whether these defects in placental vascularization were associated with altered placental oxygenation, we quantified levels of CAIX, a functional marker of placental hypoxia, in control and cannabis-exposed placentae. We demonstrated an increasing, yet non-significant, trend in positive CAIX staining of cannabis-exposed placentae (**Fig. S2**).

**Figure 5.**
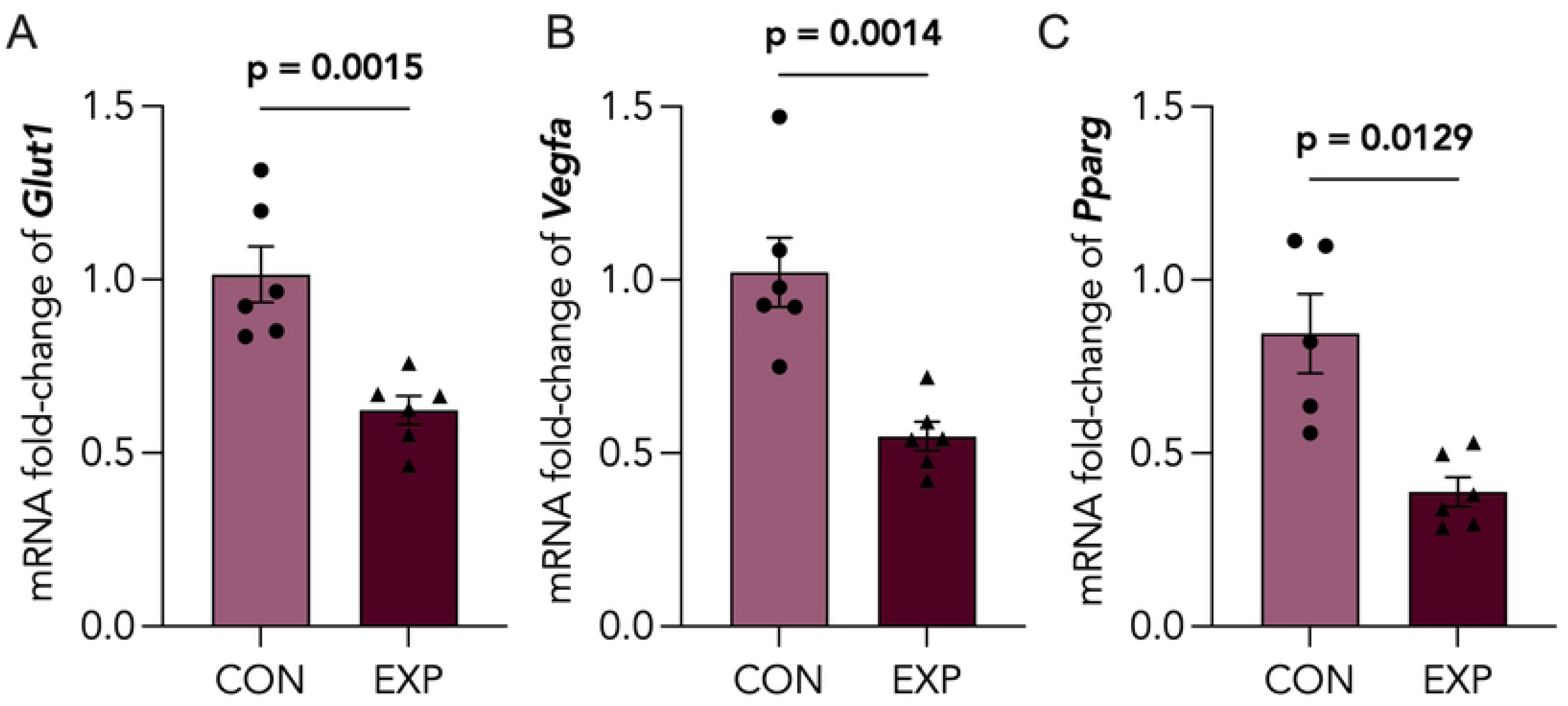
Placental nutrient transport and vascularization markers are decreased following cannabis smoke exposure. Control (CON) and cannabis-exposed (EXP) placentas were assessed for mRNA expression of (**A**) *Glut1*, (**B**) *Vegfa* and (**C**) *Pparg* via RT-qPCR, and normalized to *Actb* and *Rn18s.* Data points represent the mean ± SEM of 3-6 biological replicates. Significant differences are denoted by bolded p-values.

## Discussion

To date, few preclinical models of cannabis smoke exposure during pregnancy exist (33). In this study, we characterized a murine model for prenatal cannabis smoke exposure using an Indica-dominant hybrid strain containing 12-14% Δ9-THC and 0-2% CBD, and investigated its impacts on placental development. We quantified the total level of THC and THCA in the livers of cannabis smoke-exposed pregnant dams at E18.5 to be 135.95 ± 13.60 ng/g (Fig. 1B). Since serum THC is quickly metabolized and sequestered in lipophilic tissues (39–42), quantifying hepatic THC and THCA levels is advantageous, as it allows for more stable cannabinoid detection and assessment of hepatic cannabinoid metabolite accumulation (43). Overall, the levels THC detected in the livers of the dams are consistent with plasma THC concentrations of 45-1555 ng/ml, the reported liver and serum THC concentrations in humans who smoke cannabis (44–48).

Cannabinoids have been shown to cross the placental barrier, ultimately entering fetal tissues (49, 50). Previous studies have reported THC levels around 100 – 432 ng/g in human placental tissue and 3.9 –281.7 ng/g in the associated aborted fetuses (51), demonstrating the transplacental transfer of cannabis components. Based on our detection of THC and THCA in fetal hepatic tissue (30.84 ± 4.68 ng/g), we determined that our model of gestational cannabis smoke exposure resulted in approximately 22% of maternal THC transmission to the fetus (Fig. 1B). This is consistent with the reported range of 10-30% fetal THC transmission in preclinical models of gestational cannabinoid exposure (24, 49, 50, 52).

As the physical barrier separating maternal and fetal tissues, the placenta provides structural protection to the growing fetus through selective permeability to environmental toxicants (53). This protection is attributed, in part, to its expression of active transporters, such as P-glycoproteins, and phase I and II drug-metabolizing enzymes, including cytochrome P450 enzymes (CYPs) (54). *Cyp1a1* is an enzyme selectively induced by PAHs and aryl-containing compounds found in smoke, which functions to metabolize combustion byproducts. *Cyp1a1* is expressed by the placenta of both humans and rodents throughout gestation, suggesting its key role during placentation and pregnancy. In fact, the CYP1A1 enzyme regulates key pregnancy hormones, such as pregnenolone and estrogen intermediates, which play vital roles in the proper development and function of the placenta (55, 56). We demonstrated a significant induction of placental *Cyp1a1* mRNA following prenatal cannabis smoke exposure in mice (Fig. 1C), indicating that cannabis smoke indeed reaches placental tissue and activates xenobiotic metabolism in this model. This finding had only been previously demonstrated with prenatal tobacco exposure in vivo, whereas non-smoke exposed placentae show relatively negligible *Cyp1a1* expression both in clinical and preclinical studies (57). As placental *Cyp1a1* overexpression is associated with poor pregnancy and fetal outcomes, including intrauterine growth restriction (IUGR) (56) and growth retardation (58), our findings may offer implications for cannabis smoke-induced complications in pregnancy progression and fetal development. By contrast, cannabinoids, such as THC, have only been shown to inhibit *Cyp1a1* activity in vitro, without affecting its expression (59, 60).

Prenatal cannabis use has been associated with adverse pregnancy outcomes; however, there remains conflicting evidence in the literature regarding its impact, with some studies reporting no impacts to pregnancy outcomes (61). With 44% of pregnant people reporting co-use of tobacco, alcohol or cannabis (62), part of the discrepancy in this literature may be explained by inconsistent exclusion criteria surrounding polysubstance use. Thus, components in tobacco and alcohol may confound the impacts of cannabis exposure alone (63, 64) and increase ambiguity. As mentioned, smoked cannabis remains the most popular method of cannabis consumption, both amongst pregnant users (9, 10) and non-pregnant users (65). Despite this, the majority of preclinical studies have primarily employed single-component models of prenatal cannabinoid exposure, primarily administering THC or CBD via intraperitoneal (*i.p.*) injections (15, 16, 18, 66, 67). To better understand the effects of cannabis smoke exposure on pregnancy outcomes we assessed litter size and birthweight and found no significant differences between groups. (Fig. 2). However, the cannabis exposed animals did have a reduced placental weight, which is consistent with other clinical and preclinical studies that have reported altered placental weights or adverse pregnancy outcomes related to placental dysfunction following maternal cannabis use (68). Perhaps most concerning is the fact that we observed a significant reduction in brain:body weight ratio at birth. Importantly, this finding challenges existing work demonstrating THC-induced (*i.p* injections) symmetrical fetal growth restriction (FGR), evidenced by decreased fetal weight coupled to decreased brain- and liver-to-body weight ratios at birth (16). Notably, the clinical association between in utero cannabis smoke exposure and low birthweight remain unclear (69). Indeed, clinical studies investigating the relationship between prenatal exposures, like opioids (70) and tobacco (71), and brain size in infants, found decreased fetal brain volumes even after adjusting for birth weight, suggesting that certain environmental exposures can selectively reduce brain size. Similar to opioids and tobacco (72, 73), existing studies have highlighted the potential of cannabis smoke and its components to directly alter neurogenesis (74) and gliogenesis (75), the latter being the primary neurodevelopmental process in late gestation and early postnatal life. Overall, cannabis has been shown to cross the blood-brain barrier and accumulate in the fetal brains (24, 52); thus, it is plausible that the decreased brain-to-body weight ratio is the result of cannabis-induced dysregulation in fetal brain growth.

The structural adaptations in placental zonation that accompany changes to placental mass may allow the placenta to sustain fetal nutrient demands amid environmental pressures, thus maintaining fetal growth (76). As such, we quantified the areas of junctional and labyrinth zones from PAS-stained placental sections, in both control and cannabis-exposed placentae. In conjunction with reduced placental weights, we observed decreased junctional zone area (Fig. 3B) and increased labyrinth zone areas (Fig. 3C) in E18.5 cannabis-exposed placentae. Likewise, Chang et al. demonstrated decreased placental junctional zone area following prenatal Δ9-THC injections in mice, suggesting that Δ9-THC may be driving the disruption of junctional zone development in whole cannabis smoke (15). Interestingly, in placentae of tobacco smokers, increased villous hyperplasia was observed, ultimately mimicking the increased labyrinth zone area observed in our study, which could suggest that smoke components may also be contributing to changes in placental zonation (77).

By contrast, prenatal CBD injections did not show alterations to placental zonation in a rodent model, highlighting the differential impacts of distinct cannabinoids in cannabis on placentation (18). During early murine placentation, junctional zone formation originates from the spongiotrophoblast and ectoplacental cone trophoblasts around E7.5-E8.5 (78) and is well-established at the onset of labyrinth zone development by E12.5. Nevertheless, both zones achieve peak functionality around a similar developmental window. Therefore, it is highly plausible that delayed labyrinth zone formation may compensate for deficits that arise in junctional zone formation.

Based on these structural changes, we assessed the levels of key placental developmental markers in murine pregnancy related to junctional- or labyrinth-specific cell types, such as transcription factor AP- 2 gamma (*Tfap2c*), trophoblast specific protein alpha (*Tpbpa*) and pro-cadherin 12 (*Pcdh12*). Of interest in the context of junctional zone development are *Pcdh12* and *Tpbpa*, which are expressed primarily by glycogen-committed and spongiotrophoblast cells, respectively. We observed decreased gene expression of both *Pcdh12* (Fig. 3F) and *Tpbpa* (Fig. 3E) in cannabis-exposed placentae – an observation which may underlie the apparent reduction in junctional zone area. Furthermore, *Tfap2c* is the most highly expressed of the AP-2 family in the placenta and plays a critical role in early placental development by regulating trophoblast stem cell fate and later supports placentation by promoting invasive trophoblast development essential for implantation (79–82). We found the transcript levels of *Tfap2c* to be significantly downregulated in cannabis-exposed placentae (Fig. 3D), indicating impaired trophoblast differentiation. Interestingly, prior studies have shown prenatal 3 mg/kg CBD injections to have no effect on *Pcdh12* expression yet significantly reduced *Tfap2c* and *Tpbpa* levels in E19.5 rat placentae (18). Taken together, our findings suggest that cannabis smoke may lead to a broader downregulation of placental developmental markers, indicating the involvement of constituents in cannabis smoke beyond cannabinoids alone.

Placental endocrine function is crucial for maintaining placental and fetal growth, especially in the context of brain development (83). Thus, we assessed levels of placental lactogen 2 (*Pl2)*, a peptide hormone produced by the trophoblast giant cells from mid-to-late gestation (84–86), which is implicated in human pathologies like obesity and diabetes, and has been shown to regulate placental growth by stimulating insulin-like growth factors (IGFs) (87). In addition, we assessed insulin-like growth factor receptor 1 (*Igf1r*), which binds insulin-like growth factor (*Igf1*), a key placental growth hormone produced by the placenta that regulates fetal brain growth (88). We observed decreased transcript levels of *Pl2* (Fig. 4A), as well as downregulated *Igf1r* (Fig. 4B), which may be reflective of increased circulating levels of Igf1 when considered in the context of ligand-induced receptor downregulation. Furthermore, as *Igf1r* is almost exclusively expressed in the labyrinthine layer of the murine placenta (89), which was relatively enlarged following cannabis exposure, the downregulation of this receptor is likely mediated by receptor overactivation (90). Further investigations into IGF levels following cannabis exposure may provide insights into the potential dysregulation of the IGF axis and the implications for fetal growth.

The hallmarks of placental insufficiency, broadly defined as the inadequate functioning of the placenta, are functionally observed as an insufficient nutrient and oxygen supply to the fetus, caused by poor vascularization and placental hypoxia (91, 92). Therefore, we assessed a markers of nutrient transporters (glucose transporter 1; *Glut1*) and angiogenic signals (vascular endothelial growth factor A, *Vegfa*; peroxisome proliferator-activated receptor gamma, *Pparg*) and found all transcript levels to be significantly decreased following cannabis smoke exposure (Fig. 5), supporting the possibility that cannabis smoke exposure during pregnancy results in impaired vascularization and nutrient transport. Considering there were no observable differences in offspring weights, but rather reduced placental weight at E18.5 and brain-to-body weight ratios at PND1, our findings may suggest placental insufficiency overall. Interestingly, when rhesus macaques were given daily THC edibles pre-conception and throughout gestation, they found decreased placental perfusion and fetal oxygen availability, with no changes in placental weight or fetal weights, suggesting that cannabinoids found in cannabis may contribute to the etiology of placental insufficiency (17). Existing preclinical studies of maternal hypothyroidism have revealed decreased expression of placental *Glut1* and increased labyrinth zone size (76), highlighting a potential compensatory mechanism whereby the labyrinth zone is increased to accommodate reduced nutrient transport caused by maternal stressors. This structural adaptation represents one of many placental compensatory mechanisms to combat environmental challenges and preserve fetal growth (93–97).

Given that cannabis smoking remains the most popular mode of cannabis consumption in pregnancy, our model offers the ability to address the holistic effects of gestational cannabis use. This study is one of the first to model the consequences of cannabis smoke exposure during pregnancy on placental development. Our findings, consistent with prior work comparing inhalation and injection models of prenatal cannabis exposure (23, 52), highlight distinct effects of cannabis smoke on maternal and fetal outcomes—likely influenced by differences in metabolism and bioavailability associated with the route of administration, as well as by the presence of other cannabis constituents and combustion byproducts.

## Contributions

Conceptualization, T.P., C.M., A.H., E.M.R., S.R.; methodology, T.P., E.M.R., S.R.; formal analysis. T.P., C.M, S.R.; supervision, S.R. and E.M.R.; investigation, T.P., M.S., A.M., J.K., G.L., L.J., C.L., A.M.L., T.T., G.T., S.R., E.M.R.; resources, S.R.; writing—original draft preparation, T.P., C.M., A.M., S.R.; writing—review and editing, T.P., M.S., A.M., C.M., G.L., J.K., A.M.L., T.T., L.J., C.L., A.H., G.T., J.P., E.M.R. and S.R.; funding acquisition, S.R. All authors have read and agreed to the published version of the manuscript.

## Acknowledgements

The authors acknowledge the assistance of undergraduate students, Ashwini Pugazhendhi, Kavina Uthayakumaran, Jasjeet Chhoker, Mariyam Niaz, Danny Nguyen, Maya Bozzo-Rey, Jerron Zhang, Saundarai Bhanot, Mann Badami and Aaron Thambiahpillay during animal surgeries. The authors additionally acknowledge Mary Jo. Smith from the Histology Core, McMaster Immunology Research Center (MIRC) for PAS staining.

## Conflicts of Interest

The authors declare no conflicts of interest.

## Supporting Information

**Figure S1.** Maternal body weights and food intake remain unchanged at gestational days 8, 12 and 16 following prenatal cannabis smoke exposure. Control (CON) and cannabis smoke-exposed (EXP) dams weights and food weights were measured every four days. (**A**) Delta maternal weight gain and (**B**) food consumption (g) was quantified during pregnancy. (**C**) Estimated maternal daily energy intake based on the average food consumption over each four-day period per dam, according to the energy per gram of Teklad Global 19% Protein Extruded Diet (3.3kcal/g). Significant differences are denoted by bolded p-values.

**Figure S2.** **Placental carbonic anhydrase expression is not significantly increased following prenatal cannabis smoke exposure**. Representative images of whole E18.5 placentas from (**A**) control (CON; n=4) and (**B**) cannabis smoke-exposed (EXP; n=7) dams that were sectioned (5µm) and subsequently assessed for carbonic anhydrase (CAIX) expression with a polyclonal goat anti-carbonic anhydrase antibody (R&D Systems, AF2344) via immunohistochemistry. Magnification 400x, scale bars = 100µm. (**C**) Percent immunopositive area of CAIX staining from CON and EXP placentas at E18.5. Data are represented as a bar graph with each biological sample containing the average of CAIX quantification from two images per placenta. Statistical analyses included a Welch’s t-test. Significant differences are denoted by bolded p-values. *Jz*, junctional zone; *Lz*, labyrinth zone.

**Table S1.** A list of antibodies used for immunohistochemical analyses. Table S2. A list of mouse primer sequences used for RT-qPCR.

## Notes

### Competing Interest Statement

The authors have declared no competing interest.

